# Predicted Cellular Immunity Population Coverage Gaps for SARS-CoV-2 Subunit Vaccines and their Augmentation by Compact Peptide Sets

**DOI:** 10.1101/2020.08.04.200691

**Authors:** Ge Liu, Brandon Carter, David K. Gifford

## Abstract

Subunit vaccines induce immunity to a pathogen by presenting a component of the pathogen and thus inherently limit the representation of pathogen peptides for cellular immunity based memory. We find that SARS-CoV-2 subunit peptides may not be robustly displayed by the Major Histocompatibility Complex (MHC) molecules in certain individuals. We introduce an augmentation strategy for subunit vaccines that adds a small number of SARS-CoV-2 peptides to a vaccine to improve the population coverage of pathogen peptide display. Our population coverage estimates integrate clinical data on peptide immunogenicity in convalescent COVID-19 patients and machine learning predictions. We evaluate the population coverage of 9 different subunits of SARS-CoV-2, including 5 functional domains and 4 full proteins, and augment each of them to fill a predicted coverage gap.

## Introduction

All reported current efforts for COVID-19 vaccine design that are part of the United States Government’s Operation Warp Speed use variants of the spike subunit of SARS-CoV-2 to induce immune memory (Table S6). Subunit vaccines seek to reduce the safety risks of attenuated or inactivated pathogen vaccines by optimizing the portion of a pathogen that is necessary to produce durable immune memory (Moyle and Toth, 2013). Suggested coronavirus subunit vaccine components include the spike (S) protein, the receptor binding domain (RBD) of S, the S1 domain of S, the S2 domain of S, the nucleocapsid (N), the membrane (M), the envelope (E), the N-terminal domain (NTD) of S, and the fusion peptide (FP) of S (Wang et al., 2020; Yu et al., 2020; Dai et al., 2020). Subunit vaccines have been enabled by our ability to engineer and express pathogen surface components that retain their three-dimensional structure to induce neutralizing antibodies and a corresponding B cell memory. However, the production of durable immune memory rests in part upon help from T cells, which get their cues from peptides displayed by Human Leukocyte Antigen (HLA) molecules encoded by the Major Histocompatibility Complex (MHC) of genes. Since a subunit vaccine does not fully represent a pathogen, vaccine excluded pathogen peptides will not be observed during vaccination by an individual’s T cells.

We find that proposed SARS-CoV-2 subunit vaccines exhibit population coverage gaps in their ability to generate a robust number of predicted peptide-HLA hits in every individual. A *peptide-HLA hit* is the potential immunogenic display of a peptide by a single HLA allele. Subunit vaccine-based simulation of a T cell response is limited because of their limited representation of pathogen peptides, and the preferences of each individual’s HLA molecules for the peptides they will bind and display. Since HLA loci exhibit linkage disequilibrium, we use the frequencies of population haplotypes in our coverage computations. Each haplotype describes the joint appearance of HLA alleles. Cytotoxic CD8^+^ T cells observe peptides displayed by molecules encoded by an individual’s classical class I loci (HLA-A, HLA-B, and HLA-C), and helper CD4^+^ T cells observe peptides displayed by molecules encoded by an individual’s classical class II loci (HLA-DR, HLA-DQ, and HLA-DP).

## Results

We model peptide-HLA immunogenicity by combining data from convalescent COVID-19 patients as measured by the Multiplexed Identification of T cell Receptor Antigen specificity (MIRA) assay with machine learning predictions (Snyder et al., 2020; Klinger et al., 2015). The display of a peptide by an HLA molecule is necessary, but not sufficient, for the peptide to be immunogenic and cause T cell activation and expansion. The combined model predicts which HLA molecule displayed a peptide that was observed to be immunogenic in a MIRA experiment, and uses machine learning predictions of peptide display for HLA alleles not observed or peptides not tested in MIRA data (STAR Methods). We use this combined model of peptide immunogenicity to compute our estimates of vaccine population coverage and to propose augmentation peptides to close population coverage gaps.

We use two different candidate sets of peptides for subunit augmentation and de novo vaccine design: known immunogenic peptides, and all possible peptides. First, we exclusively use the set of peptides that were observed to be immunogenic in the MIRA assay (Snyder et al., 2020; Klinger et al., 2015). Second, we utilize peptides from the SARS-CoV-2 genome that have a mutation rate *<* 0.001 and a zero glycosylation probability predicted by NetNGlyc (Gupta et al., 2004) (STAR Methods).

For our vaccine coverage predictions we use previously reported estimates of HLA haplotype frequencies from Liu et al. (2020) for HLA-A, HLA-B, and HLA-C (classical class I) and HLA-DR, HLA-DQ, and HLA-DP (classical class II) to score the number of peptide-HLA hits observed for various subunits of SARS-CoV-2 with and without additional augmentation peptides. We also evaluate subunit population coverage for MHC class I using HLA haplotype frequencies from Gragert et al. (2013). (STAR Methods)

### SARS-CoV-2 subunit population coverage analysis

We first used our model of peptide immunogenicity to compute a baseline of the predicted number of peptide-HLA hits that would result from an infection by the SARS-CoV-2 virus using the HLA haplotype frequencies from Liu et al. (2020). For this task we extracted all peptides of length 8–10 (MHC class I) and 13–25 (MHC class II) inclusive from the SARS-CoV-2 proteome (STAR Methods).

We predict SARS-CoV-2 will have 318 (White), 307 (Black), and 391 (Asian) peptide-HLA hits for MHC class I on average in the respective self-reporting human populations. For an MHC class II redundant sampling we predict SARS-CoV-2 will have 5180 (White), 3871 (Black), and 2070 (Asian) peptide-MHC hits. Thus the average number of predicted SARS-CoV-2 peptide-HLA hits for MHC class I is 338 and for MHC class II 3707.

We found that all subunits of SARS-CoV-2 have gaps in their predicted human population coverage for robust peptide MHC display using EvalVax (STAR Methods). We computed the predicted uncovered population percentage of the SARS-CoV-2 subunits S, S1, S2, RBD, and NTD as a function of the minimum required predicted number of peptide-HLA hits displayed by an individual (Figure 1). An individual is *uncovered* if they are not predicted to have a specified number of peptide-HLA hits. Results for the FP, M, N, and E subunits are shown in Figure S1. We observe a negative correlation between subunit size and predicted population gap (Pearson *r* = −0.65 for MHC class I and Pearson *r* = −0.63 for MHC class II, Figure S2). The predicted fraction of the uncovered population was greatest for the smallest subunits, which is a direct consequence of their elimination of the largest number of pathogen peptides. For example, of the S protein subunits RBD is the smallest, and has the largest predicted population coverage gap. While the significance of the reduced immune footprint of subunit vaccines remains to be fully elucidated, when no or very few peptide-HLA hits are predicted a corresponding reduction in T cell activation, expansion, and memory function would be expected. Based on our prediction, the receptor binding domain (RBD) subunit had no MHC class II peptides displayed in 15.12% of the population (averaged across Asian, Black, and White self-reporting individuals). We note that the uncovered population of RBD with no predicted display of MHC class II peptides ranges from 0.811% for the population self-reporting as White, to a high of 37.287% for the population self-reporting as Asian. The high uncovered population in the Asian population is caused by the HLA haplotype frequencies in the Asian population. Thus, clinical trials need to carefully consider ancestry in their study designs to ensure that efficacy is measured across an appropriate population. For the RBD subunit, 26.357% of the population had fewer than six MHC class II peptide-HLA hits. For RBD MHC class I, the coverage gap is 1.163% for no hits and 38.186% for fewer than six hits. EvalVax predicted that on average for an S subunit vaccine the uncovered population would be 0.001% (class I) and 0.721% (class II) for no display, and 0.642% (class I) and 3.610% (class II) for fewer than 6 peptide-HLA hits.

**Figure 1:**
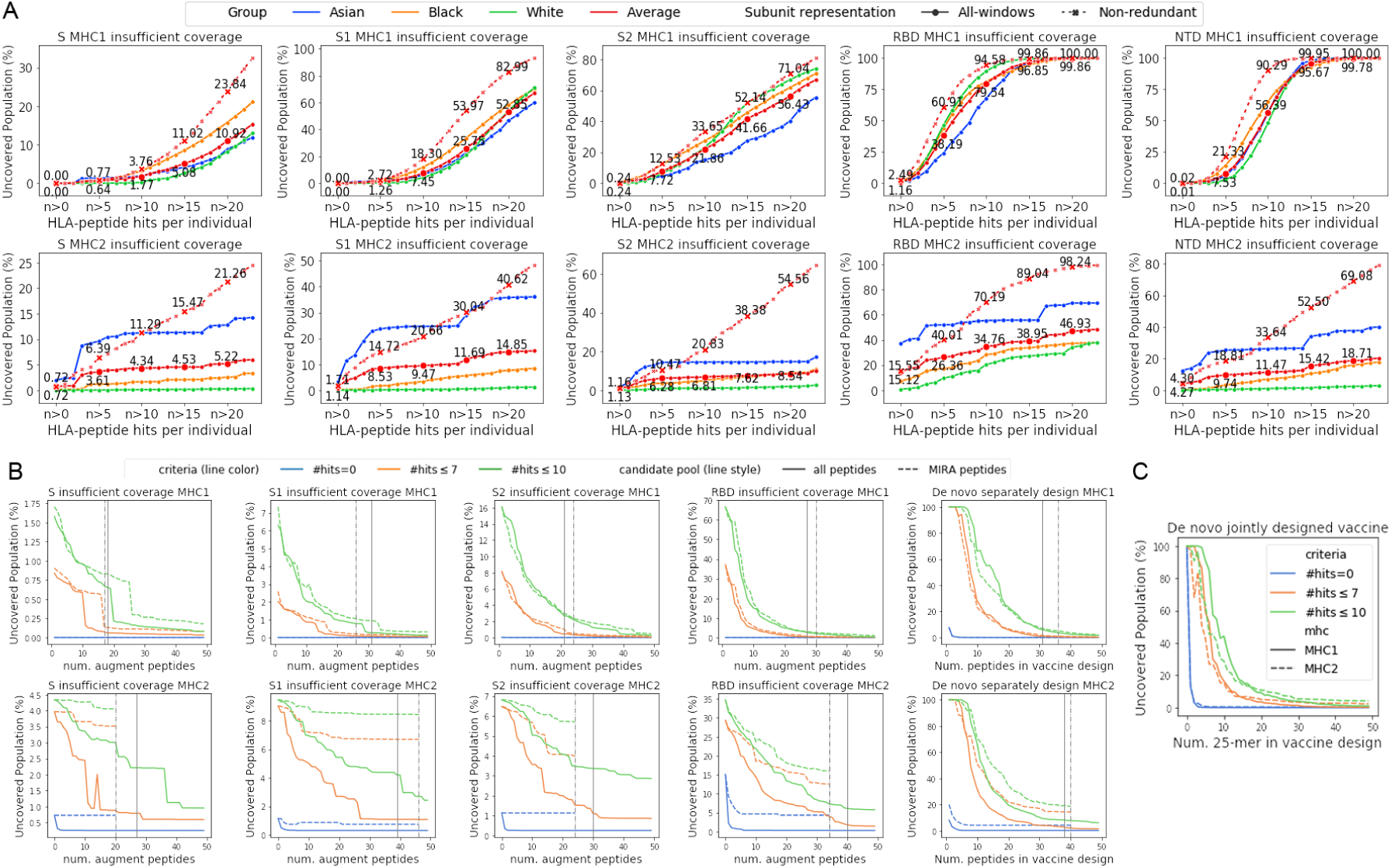
Predicted human population coverage gaps and improvement with proposed vaccines. (A) Predicted uncovered percentage of populations as a function of the minimum number of peptide-HLA hits in an individual. Annotated percentages are the average across populations self-reporting as Asian, Black, and White. A redundant sampling of peptides is depicted by solid lines for populations self-reporting as Asian, Black, and White as well their average. A non-redundant sampling of peptides is depicted by dotted lines. (B) Predicted uncovered percentage of the population for a subunit plus augmentation peptides or for a subunit free design, MHC class I (top row) and class II (bottom row). (C) Uncovered population for a joint class I and class II de novo vaccine design that does not include a subunit. Dotted graph lines in (B) utilize only MIRA validated peptides. In (B) vertical lines show the peptide count used to evaluate Table S1, dotted lines are MIRA peptides only.

We found that predicted subunit coverage gaps for MHC class I were largely consistent when we utilized HLA haplotype frequencies from Gragert et al. (2013) (Figure S5) (STAR Methods).

### SARS-CoV-2 subunit augmentation with peptide sets for MHC class I and II

We used Optivax-Robust to compute separate MHC class I and II augmentation sets of SARS-CoV-2 peptides to be combined with each subunit to maximize the predicted population coverage for a target minimum number of MHC class I and class II peptide hits in every individual (Figure 2). We used two sets of candidate peptides for these augmentation sets: (1) peptides that were found to be immunogenic in MIRA assay data, and (2) all filtered peptides from the entire SARS-Cov-2 proteome (STAR Methods). The use of peptides immunogenic in MIRA data is intended to ensure that vaccine peptides are immunogenic, while limiting population coverage by not considering other peptides that may cover rare MHC alleles. We predicted the uncovered fraction of the population as a function of MHC class I or II peptide set size for both candidate sets (Figure 1; Table S1). The computed sets of augmentation peptides were predicted to substantially reduce the populations predicted to be insufficiently covered by each subunit. Post augmentation the predicted uncovered population for RBD with no peptide-MHC hits is reduced to 0.003% (MHC class I) and 4.351% (MHC class II) with MIRA positive peptides only, and 0.0% (MHC class I) and 0.309% (MHC class II) with all filtered peptides from SARS-CoV-2. (Table S1).

**Figure 2:**
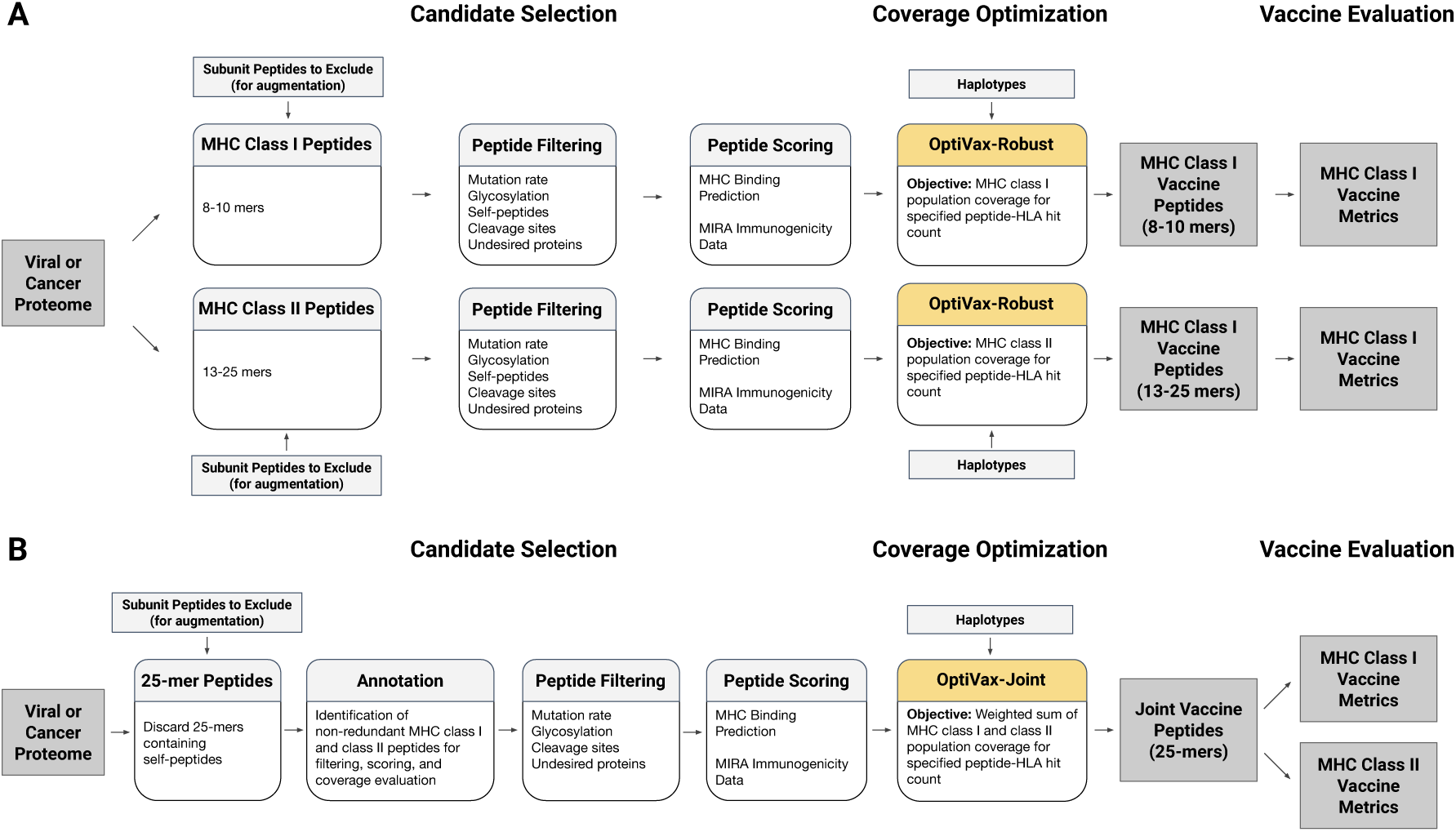
The separate and joint design methods for peptide vaccines. (A) In the separate method, windowed pathogen proteomes are filtered for acceptable peptides and MHC class I and class II vaccine designs are chosen to optimize population coverage at specified levels of peptide-HLA hits. (B) In the joint method, 25-mer pathogen peptides are annotated with their MHC class I and class II peptides, which are filtered, scored, evaluated for population coverage, and used to optimize the selection of their parent 25-mers into a joint vaccine.

### A peptide-only SARS-CoV-2 vaccine for MHC class I and II

We designed *de novo* peptide-only vaccines that did not assume an associated subunit component. We proposed either peptide vaccines that separately optimize for MHC class I and class II population coverage or a single joint peptide vaccine that optimizes MHC class I and class II coverage simultaneously (STAR methods). As candidates for vaccine inclusion we considered (1) only peptides that were found to be immunogenic in the MIRA data, and (2) all peptides from the SARS-CoV-2 proteome for separate vaccine designs. For joint vaccine design we include all peptides from the SARS-CoV-2 proteome as candidates as the MIRA positive peptides do not overlap sufficiently to do joint optimization. We explored the predicted decrease of the uncovered population as a function of peptide count and found that de novo vaccine designs are predicted to simultaneously produce a large number of predicted hits for both MHC class I and class II display (Figure 1, Table S3, Table S2). The predicted population coverage of our peptide only designs exhibit a diverse display of peptides across populations self-reporting as Black, White, and Asian (Figure S3). Peptide-only vaccine designs have been found to be effective (Herst et al., 2020).

### SARS-CoV-2 joint de novo designs are more compact than separate designs

We found that a joint de novo design that uses a single set of peptides for MHC class I and class II coverage requires fewer peptides than separate vaccines for MHC class I and class II coverage (Figure 1). With 9 jointly selected peptides more than 93% of the population was predicted to have more than 4 peptide-HLA hits. A 24 peptide joint design was predicted to produce more than 4 peptide HLA hits in more than 99% of the population (Table S2). We set a series of population coverage goals for MHC class I and MHC class II coverage with more than 7 peptide-HLA hits per individual. We considered 25 evenly spaced coverage levels between 0% and 100% coverage. We computed the total number of peptides needed to reach each set of MHC class I and MHC class II coverage goals simultaneously, where the number of MHC class I and MHC class II peptides are summed for separately designed peptide sets. We computed the total number of amino acids needed for a construct with a typical mRNA delivery platform with 10 amino acid linkers (Sahin et al., 2017) (Supplementary Information). We found joint designs reduce the total number of required amino acids and peptides required to achieve each level of population coverage (Figure S4). A single mRNA delivery construct has been demonstrated to work for both MHC class I and class II peptides (Kreiter et al., 2008; Sahin et al., 2017).

## Discussion

We augment subunit vaccines with a compact set of peptides to improve the display and immunogenicity of a vaccine on HLA class I and II molecules across a population of people. Subunit vaccines offer safety advantages over inactivated or attenuated pathogen vaccines, but their ability to fully mimic a pathogenic infection to train cellular immunity is limited. Immunity to a pathogen may rest in part upon T cell based adaptive immunity and corresponding T memory cells. We expect that a vaccine that provides a diverse display of a pathogen’s peptides will create reservoirs of CD4^+^ and CD8^+^ memory cells that will assist in establishing immunity to the pathogen. SARS-CoV-2 infection elicits a robust memory T cell response even in antibody-seronegative individuals, suggesting a T cell response is an important component of immunity to COVID-19 (Sekine et al., 2020).

We found that for SARS-CoV-2 the joint optimization of predicted MHC class I and class II pathogen peptide display achieves population coverage criteria with a more compact vaccine design than designing separate peptide sets for MHC class I and class II. Using a simpler design with shorter constructs may contribute to the effectiveness of a vaccine by providing an equivalent diversity of peptide display in a population with a less complex mixture of vaccine peptides.

Augmentation peptides can be delivered using the same vehicle as their associated subunit vaccine or they can be delivered separately. Nucleic acid based vaccines can incorporate RNA or DNA sequences that encode class I and class II augmentation peptides with desired signal sequences, linkers, and protease cleavage sites (Kreiter et al., 2008; Sahin et al., 2017) (examples in Supplementary Information, Tables S4 and S5). The peptides can be expressed as part of the subunit or separately, and can be encoded on the same or different molecules as the primary subunit. When augmentation peptides are added as a new subunit domain a vaccine designer can trade-off domain complexity for additional coverage using Figure 1B. Nucleic acid constructs carrying augmentation peptides can be delivered by injection in lipid nanoparticle particle carriers or directly (Dowdy, 2017; Wolff et al., 1990). Protein based vaccines can include independent augmentation peptides into the vaccine formulation. The delivery of independent augmentation peptides can be accomplished using nanoparticles (Herst et al., 2020).

Our computational objective function encodes the two key goals of our augmentation strategy: population coverage and the display of a highly diverse set of peptides in each individual. Our population coverage goal is ensured by optimizing predicted display coverage over population haplotype frequencies. The display of a diverse set of peptides is established by setting augmentation design goals for the number of peptides that need to be displayed by each individual.

Early results from clinical studies of subunit vaccines for SARS-CoV-2 show that some vaccine recipients did not develop positive CD8^+^ T cell responses (Jackson et al., 2020). It is difficult to fully evaluate these results because the HLA types of study participants are not provided by these early studies. Thus these study populations may not be reflective of HLA types in the general world population. The BNT162b1 RBD subunit vaccine produced a less robust CD8^+^ response than CD4^+^ response (Sahin et al., 2020), and this was also noted in the mRNA-1273 S subunit vaccine results (Jackson et al., 2020). Further clinical data is required to fully assess the T cell immunogenicity of various subunits and delivery methods. Clinical trials should select their participants to have representative HLA type distributions to test for population coverage. Future studies will need to examine the durability of immunity in individuals with minimal T cell response.

By simultaneously achieving the twin goals of coverage and diversity with peptides derived from a pathogen, we effectively compress the cellular immunologic fingerprint of a pathogen into a vaccine. To produce an antibody response, the subunit component of a vaccine can encode a three-dimensional epitope to stimulate neutralizing antibody production by B cells. Taken together, these two designed components, a pathogen subunit and its augmentation, will provide both B cell and T cell epitopes of a pathogen while permitting epitope selection to mitigate deleterious effects and improve population coverage.

All of our software and data are freely available as open source to allow others to use and extend our methods.

## Acknowledgements

This work was supported in part by Schmidt Futures, a Google Cloud Platform grant, and NIH grant R01CA218094 to D.K.G. We benefited from thoughtful comments from Michael Birnbaum, Mary Carrington, and Brooke Huisman. Ge Liu’s contribution was made prior to joining Amazon.

## Author Contributions

GL, BC, and DG contributed to problem definition and solution. GL designed and implemented the optimization procedure, with advice from BC and DG. GL, BC, and DG wrote the paper.

## Declaration of Interests

David Gifford is a founder and shareholder of ThinkTx.

## STAR Methods

### Resource Availability

#### Lead Contact

Further information and requests for resources should be directed to and will be fulfilled by the Lead Contact, David K. Gifford.

#### Materials Availability

This study did not generate new materials.

#### Data and Code Availability

All source data for generating population coverage curves have been deposited and are publicly available at https://github.com/gifford-lab/optivax. The peptide scoring predictions and processed haplotype frequencies are available at https://www.dropbox.com/sh/v1jcin4mh7jua14/AAB7W0Y7IXtXRL8Ehlrtvft6a?dl=0. This paper analyzes existing, publicly available data. These datasets’ accession numbers are provided in the Key Resources Table. All original code and the scripts used to generate the figures reported in this paper are publicly available at https://github.com/gifford-lab/optivax.

### Method Details

#### SARS-CoV-2 proteome and candidate peptides

The SARS-CoV-2 proteome is comprised of four structural proteins (E, M, N, and S) and open reading frames (ORFs) encoding nonstructural proteins (Srinivasan et al., 2020). We obtained the SARS-CoV-2 viral proteome from GISAID (Elbe and Buckland-Merrett, 2017) sequence entry Wuhan/IPBCAMS-WH-01/2019, the first documented case, as processed and provided by Liu et al. (2020). Nextstrain (Hadfield et al., 2018) was used to identify ORFs and translate the sequence. We use sliding windows to extract all peptides of length 8–10 (MHC class I) and 13–25 (MHC class II) inclusive from the SARS-CoV-2 proteome, resulting in 29,403 peptides for MHC class I and 125,593 peptides for MHC class II.

For vaccine augmentation we use two different candidate sets: known immunogenic peptides, and all possible peptides. First, we exclusively use the set of peptides that were observed to be immunogenic in the MIRA assay (Snyder et al., 2020; Klinger et al., 2015). In this case we use the MIRA sets identified for MHC class I and II separately. Second, we use the same filtered candidate peptide set as Liu et al. (2020), in which peptides with mutation rate *>* 0.001 or non-zero glycosylation probability predicted by NetNGlyc (Gupta et al., 2004) are filtered.

MIRA provides immunogenicity data with peptide-detail data that summarizes for each individual (MIRA experiment) the peptide sets that were found to cause T cell activation. A MIRA peptide set can be a single peptide, or a group of highly related peptides that are samples from slightly offset positions in the proteome. The MIRA subject-metadata contains the HLA types for individuals. While the HLA type of an individual provides us with the candidate HLA alleles that could display a given peptide, it does not tell us which allele displayed the peptide. The MIRA data used in this study includes 119 (MHC class I) or 8 (MHC class II) convalescent HLA-typed COVID-19 patients that were queried for CD8^+^ T cell activation for 269 peptide pools (generated from 545 peptides) or CD4^+^ activation for 56 peptide pools (generated from 251 peptides). Each peptide pool contains at most 13 MHC class I peptides or up to 6 MHC class II peptides. The patient population had 110 (MHC class I) and 22 (MHC class II) HLA alleles (Snyder et al., 2020). We included all MIRA immunogenic peptides for vaccine analysis and design, as to date no peptide has been observed to cause immunopathology that exacerbates disease severity.

Since the MIRA assay does not identify the patient HLA allele that presents a peptide and does not distinguish between individual peptides in a given pool, we built a combined model of MIRA observations and machine learning predictions to model peptide immunogenicity when presented by a specific HLA. We did not use the predicted HLA restrictions from Snyder et al. (2020) Supporting Table 2 as it identified pools of peptides, and not individual peptides and is only for MHC class I. For an HLA allele that appeared in the MIRA data and peptides that were tested, a peptide was predicted to be immunogenic when displayed by that HLA allele if (1) it was immunogenic in the MIRA data in 38% (MHC class I) or 40% (MHC class II) of individuals that had the HLA allele, and (2) it was predicted to bind to the HLA allele with an affinity of at least 500 nM. We used the prevalence of immunogenic peptides across individuals as criteria (1) as it performed better than using the prevalence of TCR sequences of immunogenic peptides. Other criteria that we explored that did not perform as well are included in Table 1. The selected criteria maximized the AUROC for prediction of the MIRA data that contained both positive and negative examples of peptide pool immunogenicity for individuals with a given HLA type (Table 1). Criteria (2) allowed us to predict the specific HLA allele(s) that displayed a peptide since MIRA data provides all of the HLA alleles for a given individual and does not provide information on which allele(s) displayed a peptide. We evaluate a peptide-HLA immunogenicity model using the MIRA data, and score a MIRA pool-individual pair positive if at least one peptide in the pool is predicted by the model to be immunogenic when displayed by one of the HLAs of the individual. When computing ROC or PRC curves where a variable decision boundary is employed, the maximum score across all pool peptides and HLAs is utilized for evaluation. Our combined model of HLA specific peptide immunogenicity predictions has a precision of 0.581 and AUROC of 0.833 (MHC class I) and precision of 0.849 and AUROC of 0.923 (MHC class II) (Table 1, Figure 3).

**Table 1:** Performance of machine learning only models and combined models on predicting MIRA assay immunogenicity results. The methods in bold are used for population coverage estimation and vaccine design.

| Model | Cutoff | Predicted Positive | Precision | Recall | Accuracy | F1 score | AUC | Average Precision | Specificity | p-value (precision) | p-value (accuracy) |
| --- | --- | --- | --- | --- | --- | --- | --- | --- | --- | --- | --- |
| *— MHC Class I —* |  |  |  |  |  |  |  |  |  |  |  |
| MHCflurry-2.0 predicted affinity | 500nM | 20757 | 0.297 | 0.833 | 0.506 | 0.438 | 0.700 | 0.412 | 0.407 | 0.000 | 1.000 |
| MHCflurry-2.0 predicted affinity | 50nM | 9048 | 0.410 | 0.502 | 0.718 | 0.451 | 0.700 | 0.412 | 0.783 | 0.000 | 0.000 |
| NetMHCpan-4.0 predicted affinity | 500nM | 16612 | 0.344 | 0.771 | 0.606 | 0.475 | 0.723 | 0.425 | 0.557 | 0.000 | 1.000 |
| NetMHCpan-4.0 predicted affinity | 50nM | 8196 | 0.435 | 0.481 | 0.735 | 0.457 | 0.723 | 0.425 | 0.812 | 0.000 | 0.000 |
| NetMHCpan-4.1 BA Rank | 0.5% | 13733 | 0.367 | 0.682 | 0.655 | 0.477 | 0.711 | 0.394 | 0.647 | 0.000 | 0.000 |
| NetMHCpan-4.1 BA Rank | 2.0% | 21446 | 0.300 | 0.870 | 0.501 | 0.447 | 0.711 | 0.394 | 0.390 | 0.000 | 1.000 |
| NetMHCpan-4.1 EL Rank | 0.5% | 13569 | 0.342 | 0.628 | 0.635 | 0.443 | 0.679 | 0.362 | 0.637 | 0.000 | 1.000 |
| NetMHCpan-4.1 EL Rank | 2.0% | 21423 | 0.292 | 0.845 | 0.490 | 0.434 | 0.679 | 0.362 | 0.384 | 0.000 | 1.000 |
| NetMHCpan-4.1 predicted affinity | 500nM | 16091 | 0.349 | 0.759 | 0.617 | 0.478 | 0.725 | 0.426 | 0.574 | 0.000 | 1.000 |
| NetMHCpan-4.1 predicted affinity | 50nM | 8482 | 0.435 | 0.499 | 0.734 | 0.465 | 0.725 | 0.426 | 0.805 | 0.000 | 0.000 |
| PUFFIN predicted affinity | 500nM | 17951 | 0.325 | 0.789 | 0.573 | 0.461 | 0.713 | 0.416 | 0.508 | 0.000 | 1.000 |
| PUFFIN predicted affinity | 50nM | 7870 | 0.435 | 0.462 | 0.737 | 0.448 | 0.713 | 0.416 | 0.819 | 0.000 | 0.000 |
| PUFFIN-NetMHC-MHCflurry predicted affinity | 500nM | 15953 | 0.345 | 0.744 | 0.614 | 0.472 | 0.715 | 0.424 | 0.575 | 0.000 | 1.000 |
| **PUFFIN-NetMHC-MHCflurry predicted affinity (Our ML-only model)** | 50nM | 7416 | 0.447 | 0.448 | 0.744 | 0.448 | 0.715 | 0.424 | 0.833 | 0.000 | 0.000 |
| PUFFIN-NetMHC-MHCflurry & TCR prevalence > 0.5 | 500nM | 10638 | 0.471 | 0.678 | 0.750 | 0.556 | 0.789 | 0.491 | 0.772 | 0.000 | 0.000 |
| PUFFIN-NetMHC-MHCflurry & TCR prevalence > 0.5 | 50nM | 5867 | 0.532 | 0.422 | 0.781 | 0.471 | 0.789 | 0.491 | 0.889 | 0.000 | 0.000 |
| PUFFIN-NetMHC-MHCflurry & TCR prevalence > 1 | 500nM | 9026 | 0.500 | 0.610 | 0.769 | 0.550 | 0.772 | 0.491 | 0.817 | 0.000 | 0.000 |
| PUFFIN-NetMHC-MHCflurry & TCR prevalence > 1 | 50nM | 5100 | 0.565 | 0.389 | 0.790 | 0.461 | 0.772 | 0.491 | 0.910 | 0.000 | 0.000 |
| PUFFIN-NetMHC-MHCflurry & TCR prevalence > 1.5 | 500nM | 7987 | 0.527 | 0.568 | 0.782 | 0.547 | 0.761 | 0.491 | 0.846 | 0.000 | 0.000 |
| PUFFIN-NetMHC-MHCflurry & TCR prevalence > 1.5 | 50nM | 4480 | 0.597 | 0.361 | 0.796 | 0.450 | 0.761 | 0.491 | 0.927 | 0.000 | 0.000 |
| PUFFIN-NetMHC-MHCflurry & peptide prevalence > 0.35 | 500nM | 8239 | 0.553 | 0.615 | 0.796 | 0.582 | 0.831 | 0.536 | 0.850 | 0.000 | 0.000 |
| PUFFIN-NetMHC-MHCflurry & peptide prevalence > 0.35 | 50nM | 4888 | 0.592 | 0.391 | 0.797 | 0.471 | 0.831 | 0.536 | 0.919 | 0.000 | 0.000 |
| **PUFFIN-NetMHC-MHCflurry & peptide prevalence > 0.38 (Our combined model)** | 500nM | 7388 | 0.581 | 0.580 | 0.806 | 0.581 | <u>0.833</u> | 0.549 | 0.874 | 0.000 | 0.000 |
| PUFFIN-NetMHC-MHCflurry & peptide prevalence > 0.38 | 50nM | 4445 | 0.615 | 0.369 | 0.801 | 0.462 | 0.833 | 0.549 | 0.930 | 0.000 | 0.000 |
| PUFFIN-NetMHC-MHCflurry & peptide prevalence > 0.40 | 500nM | 7074 | 0.592 | 0.566 | 0.810 | 0.579 | 0.831 | 0.554 | 0.883 | 0.000 | 0.000 |
| PUFFIN-NetMHC-MHCflurry & peptide prevalence > 0.40 | 50nM | 4280 | 0.624 | 0.361 | 0.802 | 0.457 | 0.831 | 0.554 | 0.935 | 0.000 | 0.000 |
| *— MHC Class II —* |  |  |  |  |  |  |  |  |  |  |  |
| NetMHCIIpan-3.2 BA Rank | 10.0% | 274 | 0.599 | 0.689 | 0.589 | 0.641 | 0.619 | 0.639 | 0.476 | 0.001 | 0.000 |
| NetMHCIIpan-3.2 BA Rank | 2.0% | 84 | 0.607 | 0.214 | 0.509 | 0.317 | 0.619 | 0.639 | 0.843 | 0.000 | 0.431 |
| NetMHCIIpan-3.2 predicted affinity | 500nM | 411 | 0.567 | 0.979 | 0.592 | 0.718 | 0.622 | 0.638 | 0.152 | 0.061 | 0.000 |
| NetMHCIIpan-3.2 predicted affinity | 50nM | 53 | 0.698 | 0.155 | 0.516 | 0.254 | 0.622 | 0.638 | 0.924 | 0.000 | 0.292 |
| NetMHCIIpan-4.0 BA Rank | 10.0% | 379 | 0.586 | 0.933 | 0.614 | 0.720 | 0.696 | 0.710 | 0.252 | 0.007 | 0.000 |
| NetMHCIIpan-4.0 BA Rank | 2.0% | 165 | 0.691 | 0.479 | 0.609 | 0.566 | 0.696 | 0.710 | 0.757 | 0.000 | 0.000 |
| NetMHCIIpan-4.0 EL Rank | 10.0% | 279 | 0.649 | 0.761 | 0.654 | 0.700 | 0.726 | 0.739 | 0.533 | 0.000 | 0.000 |
| NetMHCIIpan-4.0 EL Rank | 2.0% | 72 | 0.847 | 0.256 | 0.580 | 0.394 | 0.726 | 0.739 | 0.948 | 0.000 | 0.001 |
| NetMHCIIpan-4.0 predicted affinity | 500nM | 374 | 0.586 | 0.920 | 0.612 | 0.716 | 0.701 | 0.731 | 0.262 | 0.007 | 0.000 |
| **NetMHCIIpan-4.0 predicted affinity (Our ML-only model)** | 50nM | 61 | 0.869 | 0.223 | 0.569 | 0.355 | 0.701 | 0.731 | 0.962 | 0.000 | 0.003 |
| PUFFIN predicted affinity | 500nM | 417 | 0.561 | 0.983 | 0.583 | 0.715 | 0.664 | 0.656 | 0.129 | 0.088 | 0.000 |
| PUFFIN predicted affinity | 50nM | 141 | 0.688 | 0.408 | 0.587 | 0.512 | 0.664 | 0.656 | 0.790 | 0.000 | 0.000 |
| NetMHCIIpan-4.0 predicted affinity & TCR prevalence > 0.5 | 500nM | 249 | 0.827 | 0.866 | 0.833 | 0.846 | 0.880 | 0.864 | 0.795 | 0.000 | 0.000 |
| NetMHCIIpan-4.0 predicted affinity & TCR prevalence > 0.5 | 50nM | 59 | 0.898 | 0.223 | 0.574 | 0.357 | 0.880 | 0.864 | 0.971 | 0.000 | 0.001 |
| NetMHCIIpan-4.0 predicted affinity & TCR prevalence > 1 | 500nM | 225 | 0.858 | 0.811 | 0.828 | 0.834 | 0.872 | 0.858 | 0.848 | 0.000 | 0.000 |
| NetMHCIIpan-4.0 predicted affinity & TCR prevalence > 1 | 50nM | 57 | 0.895 | 0.214 | 0.569 | 0.346 | 0.872 | 0.858 | 0.971 | 0.000 | 0.003 |
| NetMHCIIpan-4.0 predicted affinity & TCR prevalence > 1.5 | 500nM | 204 | 0.887 | 0.761 | 0.821 | 0.819 | 0.857 | 0.851 | 0.890 | 0.000 | 0.000 |
| NetMHCIIpan-4.0 predicted affinity & TCR prevalence > 1.5 | 50nM | 55 | 0.909 | 0.210 | 0.569 | 0.341 | 0.857 | 0.851 | 0.976 | 0.000 | 0.002 |
| NetMHCIIpan-4.0 predicted affinity & peptide prevalence > 0.35 | 500nM | 265 | 0.811 | 0.903 | 0.837 | 0.855 | 0.899 | 0.878 | 0.762 | 0.000 | 0.000 |
| NetMHCIIpan-4.0 predicted affinity & peptide prevalence > 0.35 | 50nM | 59 | 0.898 | 0.223 | 0.574 | 0.357 | 0.899 | 0.878 | 0.971 | 0.000 | 0.001 |
| NetMHCIIpan-4.0 predicted affinity & peptide prevalence > 0.38 | 500nM | 265 | 0.811 | 0.903 | 0.837 | 0.855 | 0.899 | 0.878 | 0.762 | 0.000 | 0.000 |
| NetMHCIIpan-4.0 predicted affinity & peptide prevalence > 0.38 | 50nM | 59 | 0.898 | 0.223 | 0.574 | 0.357 | 0.899 | 0.878 | 0.971 | 0.000 | 0.001 |
| **NetMHCIIpan-4.0 predicted affinity & peptide prevalence > 0.4 (Our combined model)** | 500nM | 251 | 0.849 | 0.895 | 0.859 | 0.871 | <u>0.923</u> | 0.903 | 0.819 | 0.000 | 0.000 |
| NetMHCIIpan-4.0 predicted affinity & peptide prevalence > 0.4 | 50nM | 54 | 0.944 | 0.214 | 0.576 | 0.349 | 0.923 | 0.903 | 0.986 | 0.000 | 0.001 |

**Figure 3:**
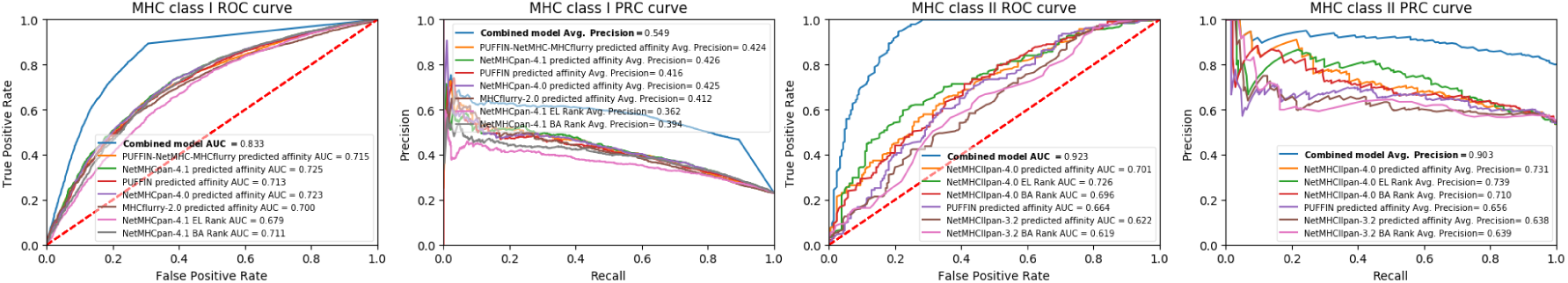
An integrated model of MIRA data and computational predictions is the best predictor of MIRA immunogenicity data. Receiver Operating Characteristic (ROC) and Precision Recall Curve (PRC) plots for predicting MIRA assay detected peptide immunogenicity using machine learning methods. The combined model curve is the performance of the integrated model of MIRA data and computational predictions.

For HLA alleles not present or peptides not tested in MIRA data we use machine learning predictions of peptide immunogenicity. We evaluated machine learning methods by their ability to predict MIRA peptides that are immunogenic in an individual based upon the HLA type of the individual (Figure 3). For a given individual we used both positive and negative sets to characterize their performance, and we prioritized precision for conservative vaccine design (Table 1). We found for MHC class I the best method utilized a 50 nM threshold from an ensemble that outputs the mean predicted binding affinity of NetMHCpan-4.0 (Jurtz et al., 2017), PUFFIN (Zeng and Gifford, 2019), and MHCflurry 2.0 (O’Donnell et al., 2020, 2018). We selected this ensemble as it is more robust to errors by a single method. For MHC class II the method we selected used a 50 nM threshold and NetMHCIIpan-4.0 (Reynisson et al., 2020b). Our machine learning predictions of HLA specific peptide display have a precision of 0.447 and AUROC of 0.715 (MHC class I) and a precision of 0.869 and AUROC of 0.701 (MHC class II) for immunogenicity (Figure 3). Other methods we explored included NetMHCpan-4.1 (Reynisson et al., 2020a) (MHC class I), PUFFIN (Zeng and Gifford, 2019) (MHC class II) and NetMHCIIpan-3.2 (Jensen et al., 2018) (MHC class II) (Table 1).

We use HLA class I and class II haplotype frequencies provided by Liu et al. (2020). HLA haplotype frequencies were generated from previously published next-generation sequencing data generated in the Carrington lab and their collaborators (Tang et al., 2012; Ramsuran et al., 2018). All of these HLA data are based upon genome sequencing that provides the highest resolving power for HLA typing. For the HLA class I locus, this dataset contains 2,138 distinct haplotypes spanning 230 HLA-A, HLA-B, and HLA-C alleles. For HLA class II, this dataset contains 1,711 distinct haplotypes spanning 280 HLA-DP, HLA-DQ, and HLA-DR alleles. Population frequencies are provided for three populations self-reporting as having White, Black, or Asian ancestry. We used these data for vaccine evaluation and design as they included the HLA-DQA and HLA-DPA/DPB alleles for MHC class II that are not present in Gragert et al. (2013).

We also predicted MHC class I population coverage using Gragert et al. (2013). For this analysis we used a combined immunogenicity model of peptide-HLA immunogenicity with an ensemble of NetMHCpan-4.0 and MHCflurry 2.0 (Jurtz et al., 2017; O’Donnell et al., 2020, 2018) for machine learning predictions.

We consider nine subunit vaccines for SARS-CoV-2: the full envelope (E), membrane (M), nucleocapsid (N), and spike (S) proteins as well as the S1, S2, receptor binding domain (RBD), N-terminal domain (NTD), and fusion peptide (FP) domains from S. The amino acid positions for each of the S protein subunits are shown in Figure 4. When evaluating these subunit vaccines we include all peptides of length 8–10 (MHC class I) and 13–25 (MHC class II) spanning the corresponding regions of the proteome.

**Figure 4:**
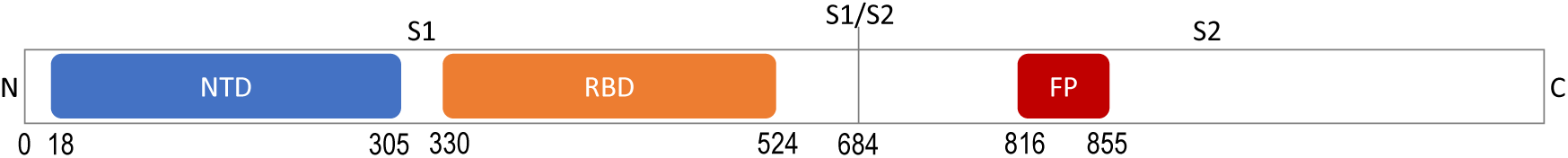
Illustration of functional domains on SARS-CoV-2 S protein.

#### EvalVax subunit vaccine evaluation

We evaluate population coverage of SARS-CoV-2 subunit vaccines using EvalVax-Robust (Liu et al., 2020). EvalVax-Robust computes population coverage of a given peptide set using the HLA haplotype frequencies in each population of individuals self-reporting as having Black, Asian, or White ancestry. Population coverage *P* (*n*) is defined as the fraction of individuals predicted to have ≥ *n* peptide-HLA hits using our model of peptide-HLA immunogencity. EvalVax-Robust computes the frequency of diploid HLA genotypes, and accounts for both homozygous and heterozygous HLA loci. We compute the average population coverage as an unweighted average of population coverage over the three populations. Insufficient coverage of ≤ *n* hits is defined as 100% − *P* (*n* + 1).

Our subunit population coverage estimates are not lowered by discarding subunit peptides as unsuitable. We consider all peptides that result from a windowing of the subunit proteome, and include the redundant peptides caused by using varying window sizes at the same proteome start position. In addition, we do not filter peptides for mutation rate or glycosylation during evaluation.

#### Design of separate MHC class I and II peptide sets to augment subunit vaccine population coverage

In the separate design method we use OptiVax-Robust (Liu et al., 2020) to augment subunit vaccines with additional peptides to produce separate sets of peptides for class I and class II augmentation (Figure 2A). The candidate peptides for vaccine inclusion are chosen from either: (1) all peptides observed to be immunogenic in a MIRA assay, or (2) all filtered peptides from the SARS-CoV-2 proteome. All filtered peptides are selected from the remaining SARS-CoV-2 proteome (all peptides except those spanning the subunit), excluding peptides that are likely to mutate (have mutation rate *>* 0.001) or have non-zero predicted probability of glycosylation. All candidate peptides considered during augmentation must be predicted to be immunogenic using our model of peptide-HLA immunogenicity.

The augmentation algorithm uses a starting peptide set which is extracted from the subunit vaccine to maximize the coverage of the subunit while removing redundant peptides resulting from overlapping sliding windows using the redundancy elimination algorithm found in Liu et al. (2020). Using a non-redundant starting peptide set ensures that augmentation does not depend upon redundant peptides for population coverage support. OptiVax-Robust performs vaccine augmentation by adding peptides to this starting set to improve the population coverage at each peptide-HLA hits cutoff *n*. At each iteration redundant peptides are removed from consideration, and redundancy is defined with an edit distance metric (Liu et al., 2020). OptiVax-Robust uses a beam search algorithm that iteratively expands the solution by one peptide and gradually optimizes population coverage from *n* = 1 to the targeting level of per-individual peptide-HLA hits (Liu et al., 2020). We use a beam size of 5 for the augmentation of subunit vaccines.

For each desired budget of augmentation peptides, OptiVax produces an augmentation set. Larger augmentation sets are not necessarily supersets of smaller augmentation sets, as the underlying combinatorial optimization problem is complex. A vaccine designer can evaluate how many peptides they wish to use to realize a predicted population coverage. For the augmentation sets in Table S1 for *n* = 7 we targeted 99.3% coverage for MHC class I augmentation and 98% coverage for MHC class II. The exceptions were S and S1, where we targeted for MHC class I 99.9% coverage (all peptides) or 99.7% (MIRA peptide only), and for class II 98.5% (all peptides) or 98% (MIRA peptides). Class II is more difficult to cover with MIRA peptides alone, and thus we accept the best coverage possible. Augmentation sets are computed starting with non-redundant subunits to avoid peptide-hit credit for windowing induced redundancies. For the evaluation of original and augmented subunit vaccines in Table S1, we provide results for all window derived subunit peptides and the non-redundant set of subunit peptides. All window peptides can include the same HLA binding epitope multiple times from its sampling by multiple windows, and thus serves as the predicted lower bound on population insufficient coverage. The non-redundant results are the predicted upper bound of population insufficient coverage.

### Design of a single set of peptides to maximize MHC class I and II population coverage

We developed the OptiVax-Joint method to produce a minimal set of 25-mer peptides to reach a target population coverage probability at a threshold of *n* predicted hits for each individual for both MHC class I and class II (Figure 2B). The 25-mer candidate peptides are produced by windowing the pathogen proteome that is not part of a selected subunit, using a window step size of 8 amino acids between candidate peptides. Each of the candidate 25-mer peptides is annotated with its non-redundant peptides of length 8–10 (MHC class I) and 13–25 (MHC class II) and the HLA alleles where they are predicted to be immunogenic. Peptide redundancy is defined with an edit distance metric for the elimination of overlapping peptides (Liu et al., 2020).

OptiVax-Joint begins with the empty set, and performs vaccine augmentation by adding candidate 25-mer peptides to this starting set to improve both MHC class I and class II population coverage at a target number of peptide-HLA hits *n*. When OptiVax-Joint is started with an empty set of peptides it produces a *de novo* peptide vaccine design without an associated subunit component. Each 25mer is scored based on its contained annotated class I and class II peptides for its improvement in the number of per-individual peptide-HLA hits (Liu et al., 2020) over the haplotypes of the target population. Contained peptides are not counted towards population coverage if they have an observed mutation rate *>* 0.001 or have a non-zero predicted probability of glycosylation. OptiVax-Joint uses a beam search algorithm that iteratively expands the solution by one 25-mer peptide and gradually optimizes population coverage from *n* = 1 peptide hit to the targeted level of per-individual peptide-HLA hits for both MHC class I and class II (Liu et al., 2020). We use a beam size of 5 for the augmentation of subunit vaccines.

For each desired budget of peptides, OptiVax-Joint produces a vaccine peptide set. Larger sets are not necessarily supersets of smaller augmentation sets, as the underlying combinatorial optimization problem is complex. A vaccine designer can evaluate how many peptides they wish to use to realize a predicted population coverage. For the joint sets in Table S1, we targeted 99% coverage at *n* = 7 for MHC class I augmentation and 97% coverage at *n* = 7 for MHC class II augmentation.

As a point of comparison, we also computed separate MHC class I and class II vaccine designs using OptiVax-Robust, using candidate sets drawn either from MIRA immunogenic peptides or all filtered peptides.

### Quantification and Statistical Analysis

Classification performance of peptide-MHC scoring models was calculated using scikit-learn (Pedregosa et al., 2011) in Python using the *sklearn.metrics.roc auc score* (AUROC), *sklearn.metrics.average_precision_score* (Average Precision), *sklearn.metrics.accuracy_score* (Accuracy), *sklearn.metrics.precision recall fscore support* (Precision, Recall and F1 score), and *sklearn.metrics.classification_report* (Sensitivity and Specificity) functions. AUROC and average precision are computed using raw predictions, and the remaining metrics are computed using binarized predictions based on the respective binding criteria. Pearson *r* correlation was computed using scipy (Virtanen et al., 2020) in Python using the *scipy.stats.pearsonr* function.

## Supplementary Information

**Figure S1:**
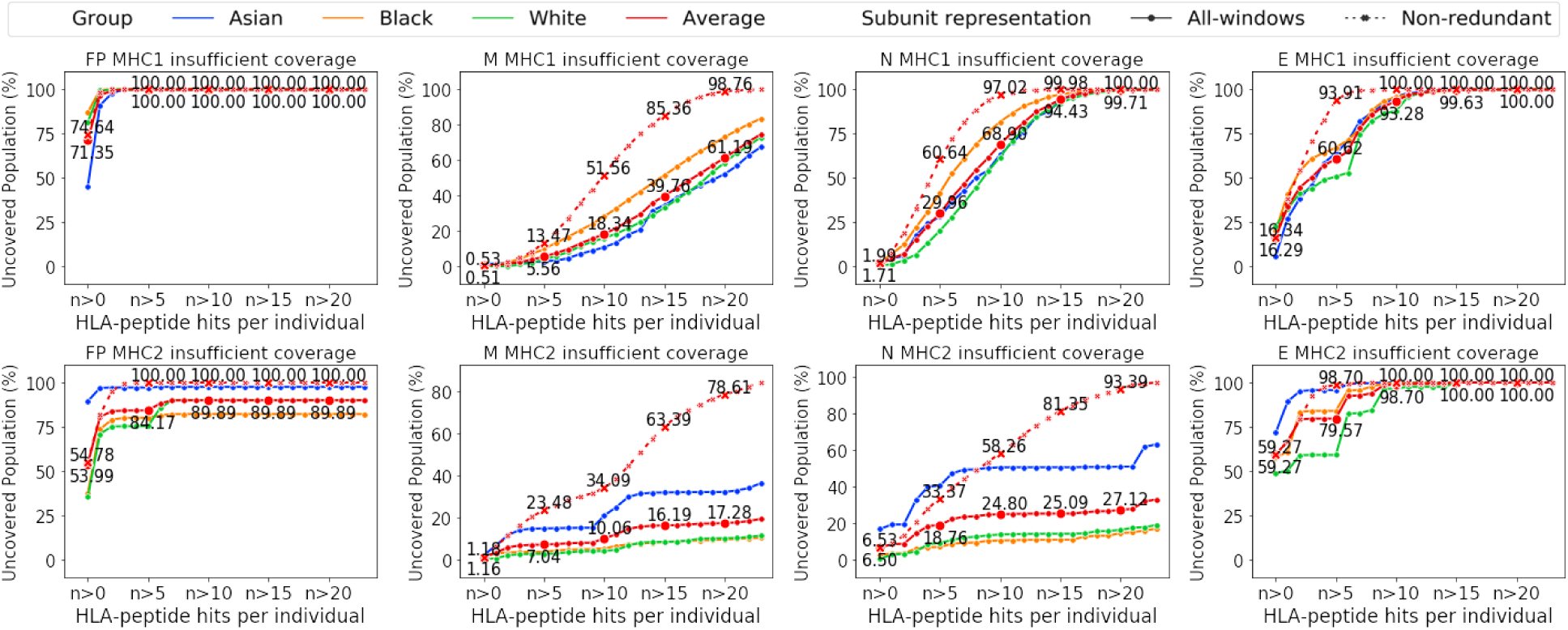
Predicted uncovered percentage of populations as a function of the minimum number of peptide-HLA hits in an individual for E, M, N protein and fusion peptide (FP). Annotated percentages are the average across populations self-reporting as Asian, Black, and White. A redundant sampling of peptides is depicted by solid lines for populations self-reporting as Asian, Black, and White as well their average. A non-redundant sampling of peptides is depicted by dotted lines.

**Figure S2:**
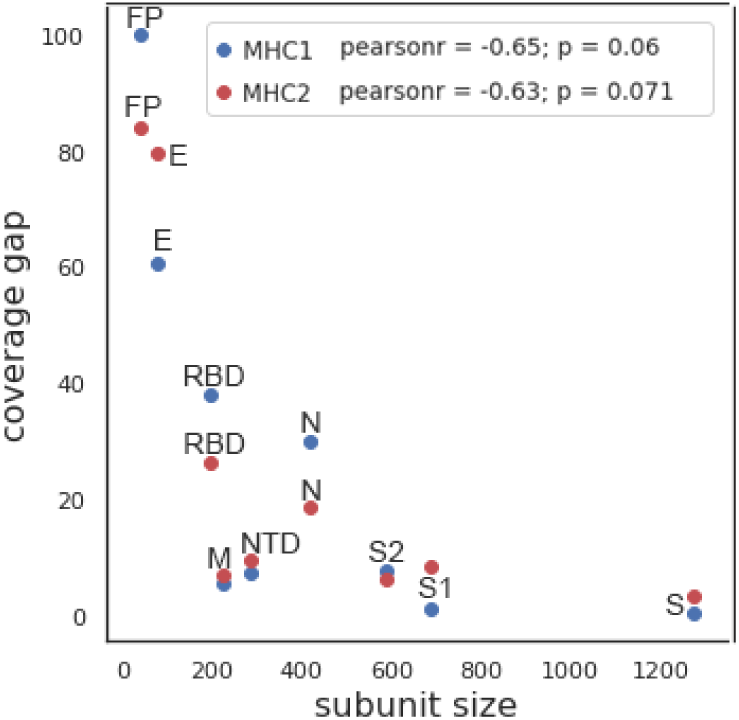
Correlation between subunit size and the predicted population gap in percent of population with less than six peptide-HLA hits per individual for MHC class I (blue) and MHC class II (red).

**Figure S3:**
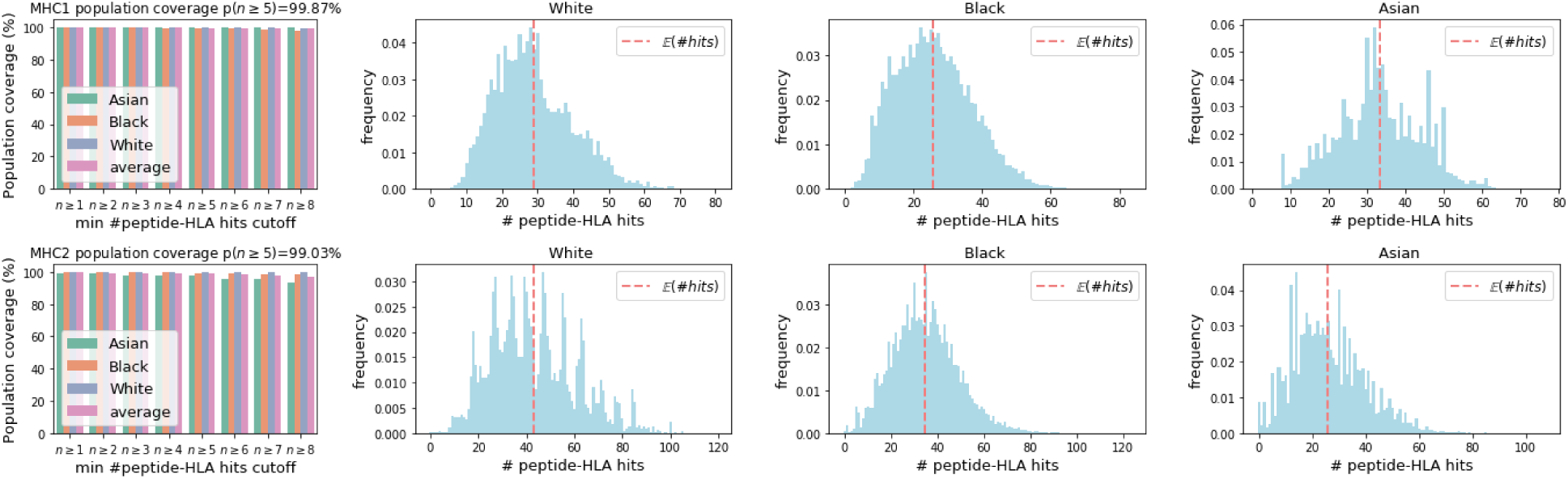
Predicted coverage in populations self-reporting as White, Black, and Asian with a peptide-only vaccine comprising 24 25-mer peptides jointly optimized for MHC class I and MHC class II coverage. The red dotted vertical line shows the expected number of hits.

**Figure S4:**
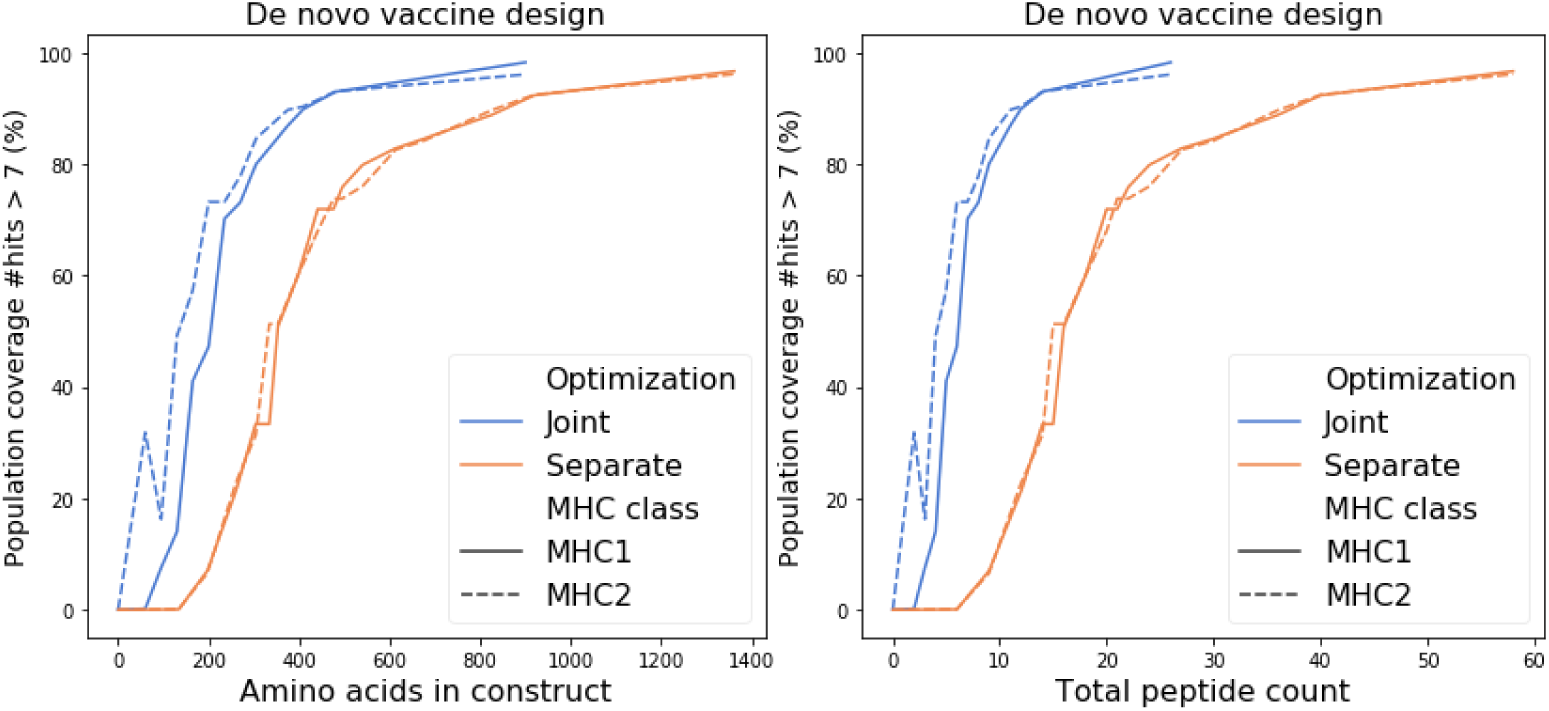
Comparison of the number of amino acids and total number of peptides used by separate and joint designs and their respective predicted population coverage with more than 7 peptide-HLA hits per individual. The predicted population coverage is shown for the peptide-only *de novo* vaccine designs.

**Figure S5:**
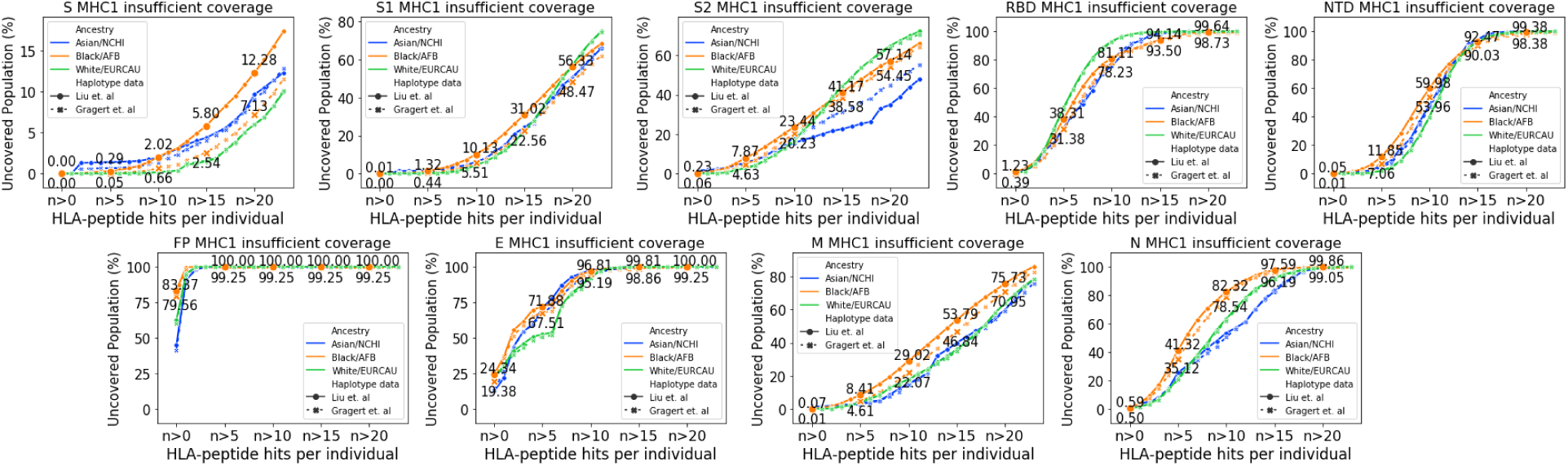
Comparison of predicted human population coverage gaps using MHC class I HLA haplotype frequencies from Gragert et al. (2013) and Liu et al. (2020). Predicted uncovered percentage of populations as a function of the minimum number of peptide-HLA hits in an individual. HLA haplotype frequencies are from Gragert et al. (2013) (dotted lines) or Liu et al. (2020) (solid lines). Annotated numeric percentages are the average population gaps across populations self-reporting as Black/AFB with haplotype frequencies from Gragert et al. (2013) and Liu et al. (2020). All data are for the redundant sampling of subunits. Peptide scoring is based upon MIRA data as elaborated by machine learning predictions by an ensemble of NetMHCpan4.0 and MHCflurry 2.0.

**Table S1:**
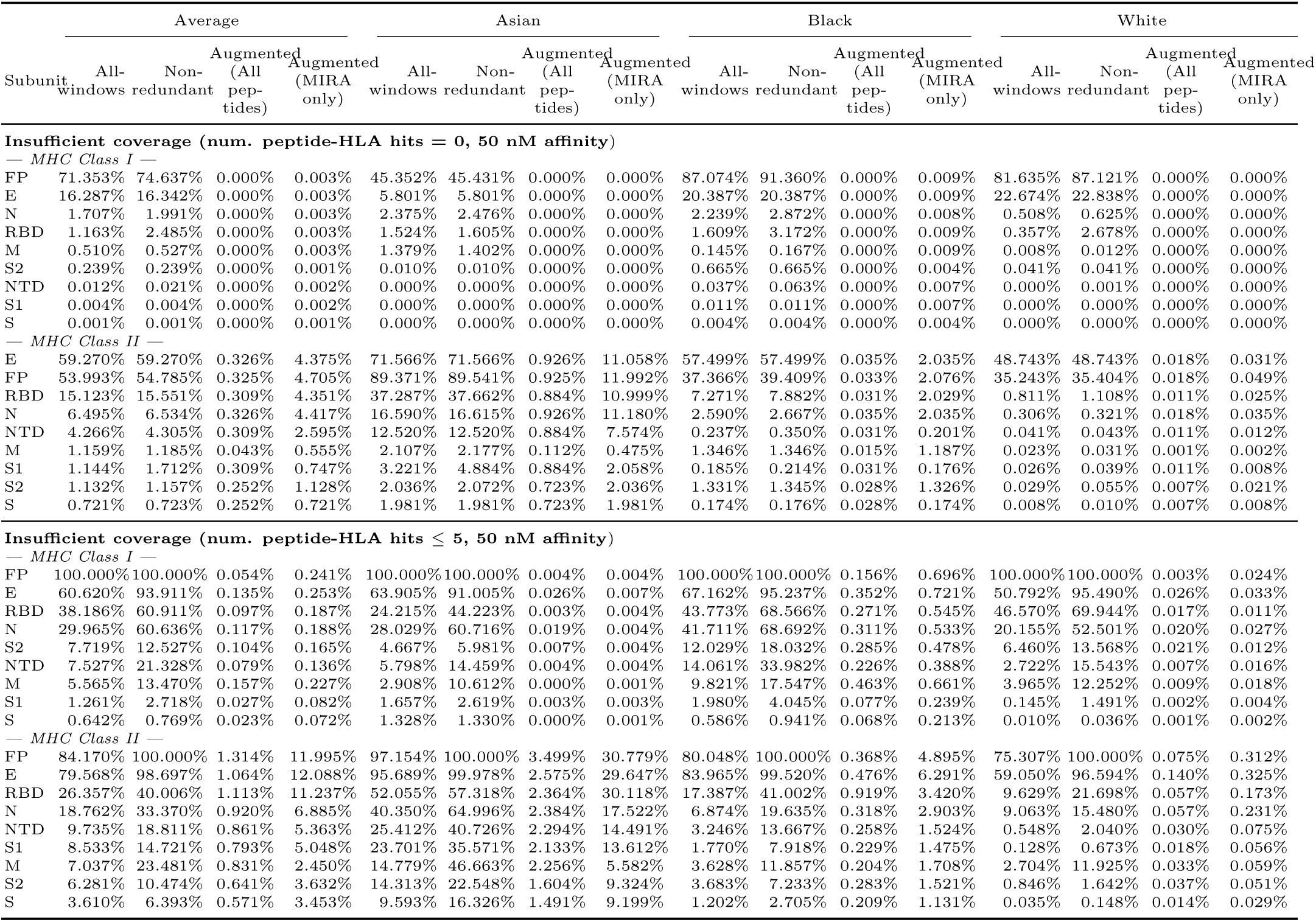
Percentage of a population that is insufficiently covered by subunit vaccines and the improvement after adding MHC class I and MHC class II augmentation peptides. Results are shown for both separate and joint designs of augmentation peptides. The list is sorted by decreasing insufficient coverage of unaugmented subunits. We chose the set with the minimal number of peptides that achieves the targeting criteria specified by OptiVax-Joint.

**Table S2:**
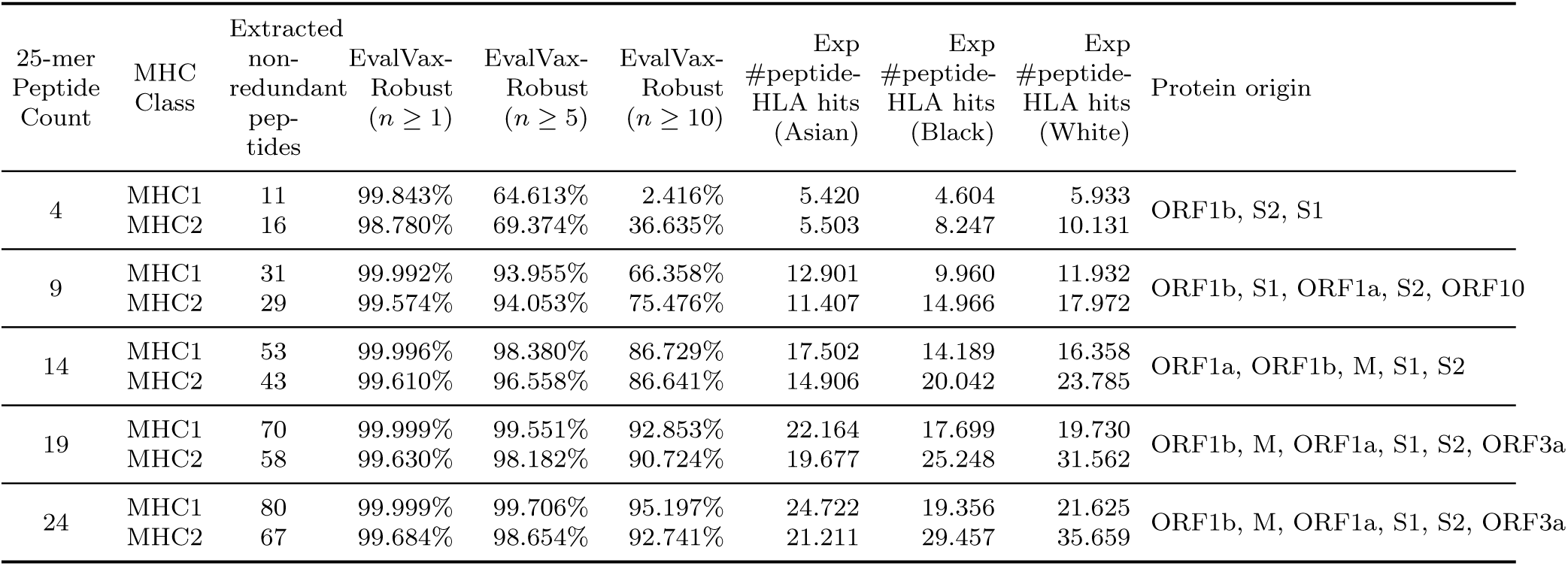
Predicted population coverage of a peptide-only vaccine jointly optimized for MHC class I and class II coverage with 4, 9, 14, 19, and 24 25-mer peptides.

**Table S3:** Predicted population coverage of peptide-only vaccines optimized separately for MHC class I and class II coverage, using either MIRA positive peptides only or all filtered peptides in SARS-CoV-2.

| Candidates | MHC<br>Class | Peptide<br>Count | EvalVax-Robust<br>($n \geq 1$) | EvalVax-Robust<br>($n \geq 5$) | EvalVax-Robust<br>($n \geq 10$) | Exp<br>#peptide-HLA hits<br>(Asian) | Exp<br>#peptide-HLA hits<br>(Black) | Exp<br>#peptide-HLA hits<br>(White) | Protein origin |
| --- | --- | --- | --- | --- | --- | --- | --- | --- | --- |
| MIRA<br>peptides | MHC1 | 36 | 99.997% | 99.832% | 96.614% | 22.987 | 20.017 | 21.957 | M, N, ORF10, ORF1a,<br>ORF1b, ORF3a, ORF7a, S1,<br>S2 |
| All peptides | MHC1 | 31 | 100.000% | 99.949% | 96.366% | 18.959 | 17.026 | 18.821 | M, N, ORF10, ORF1a,<br>ORF1b, ORF3a, S1, S2 |
| MIRA<br>peptides | MHC2 | 40 | 95.625% | 88.796% | 81.923% | 16.560 | 26.818 | 39.001 | M, N, ORF3a, ORF7a, ORF8,<br>S1, S2 |
| All peptides | MHC2 | 38 | 99.691% | 99.042% | 92.767% | 19.041 | 26.220 | 29.402 | M, N, ORF1a, ORF1b,<br>ORF3a, S1, S2 |

**Table S4:**
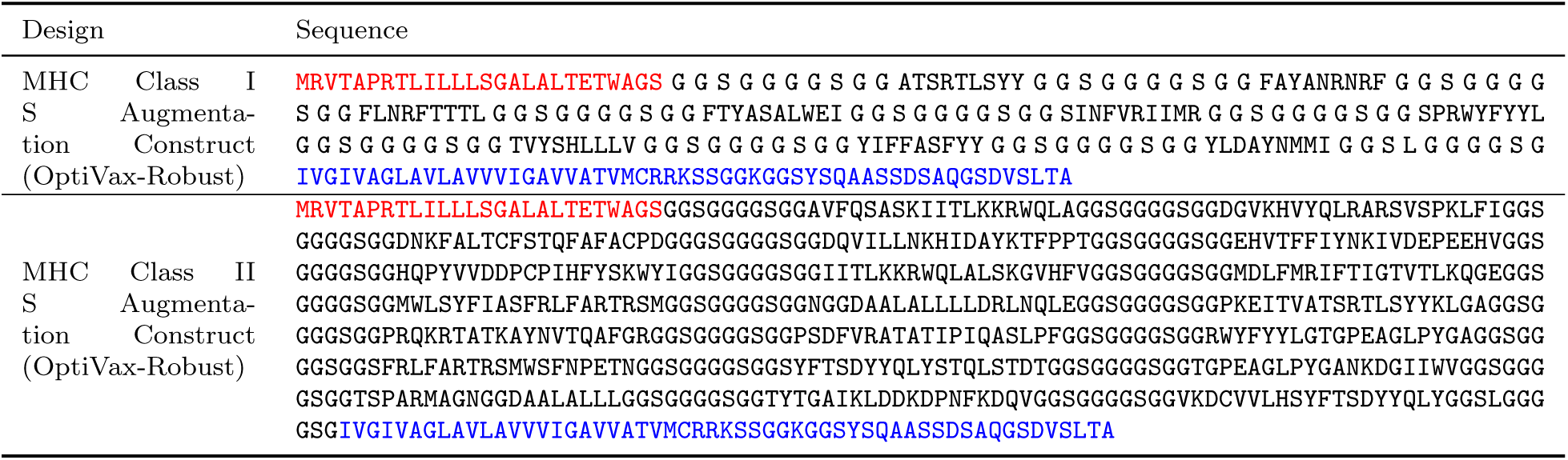
Example protein constructs for augmentations to S subunit vaccines optimized for either MHC class I or MHC class II by OptiVax-Robust. Constructs contain a secretion signal sequence (red), peptides (bold) joined by non-immunogenic glycine/serine linkers, and an MHC class I trafficking signal (blue). The augmentation peptides encoded are the same as those evaluated in Table S1. Peptides are prepended with a secretion signal sequence at the N-terminus and followed by an MHC class I trafficking signal (MITD) (Kreiter et al., 2008; Sahin et al., 2017). The MITD has been shown to route antigens to pathways for HLA class I and class II presentation (Kreiter et al., 2008). Here we combine all peptides of each MHC class into a single construct using a non-immunogenic glycine/serine linkers from Sahin et al. (2017), though it is also plausible to construct individual constructs containing single peptides with the same secretion and MITD signals as demonstrated by Kreiter et al. (2008).

**Table S5:**
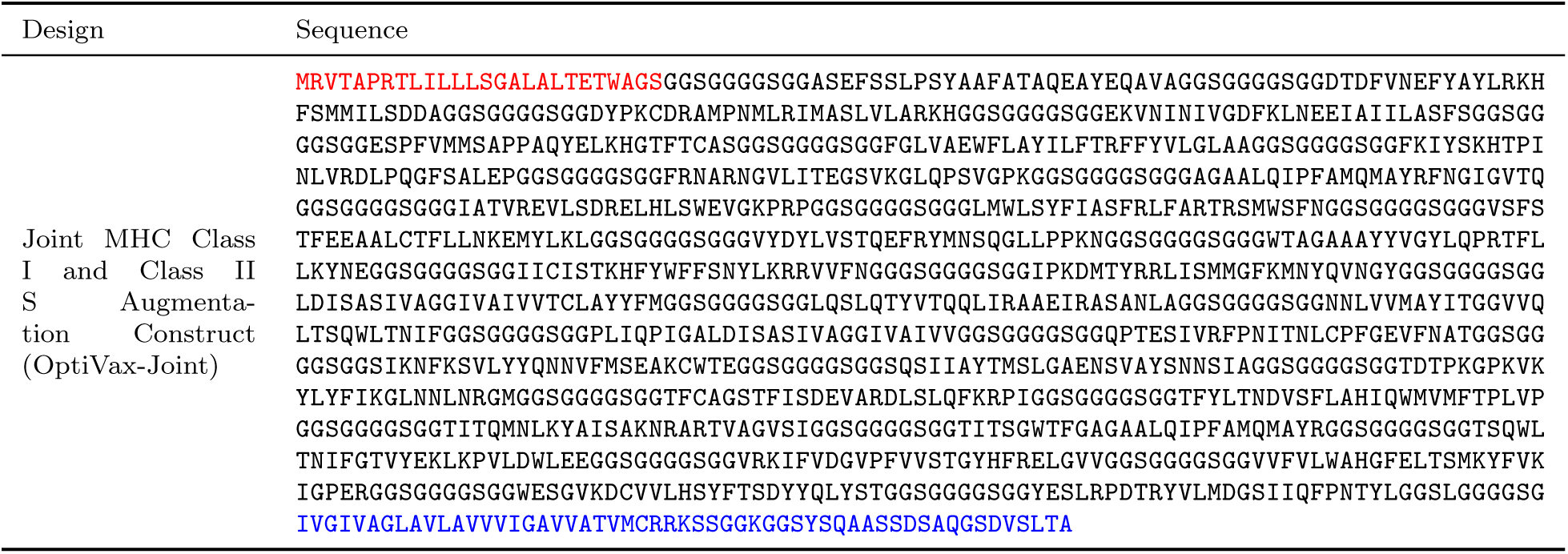
Example protein construct for augmentations to S subunit vaccines jointly optimized for both MHC class I and class II by OptiVax-Joint. Constructs contain a secretion signal sequence (red), 33 25-mer peptides (bold) joined by non-immunogenic glycine/serine linkers, and an MHC class I trafficking signal (blue). The augmentation peptides encoded are the same as those evaluated in Table S1.

**Table S6:** Overview of SARS-CoV-2 vaccines in development by Operation Warp Speed participants. See World Health Organization (2020) for additional COVID-19 candidate vaccines.

| Organization | Vaccine | Subunit | Delivery Method | Reference |
| --- | --- | --- | --- | --- |
| Janssen (Johnson & Johnson) | Ad26COVS1 | S1 | Adenoviral vector | Belgian Biosafety Server (2020) |
| AstraZeneca + University of Oxford | AZD1222 (ChAdOx1 nCoV-19) | S | Adenoviral vector | Folegatti et al. (2020) |
| Pfizer + BioNTech | BNT162b1<br>BNT162b2 | RBD<br>S | mRNA<br>mRNA | Mulligan et al. (2020)<br>BioNTech (2020) |
| Moderna | mRNA-1273 | S | mRNA | Jackson et al. (2020) |
| Merck + IAVI | — | S | VSV chimeric virus | Cohen (2020) |
| Merck + Themis | — | S | Attenuated measles virus | Cohen (2020) |
| Novavax | NVX-CoV2373 | S | Protein subunit (nanoparticle) | Tian et al. (2020) |
| Sanofi + GlaxoSmithKline | — | S | Protein subunit | World Health Organization (2020) |

Table S7: Detailed augmentation designs for optimized peptide sets in Table S1. See `AugmentationPeptides.xlsx`.

Table S8: List of uncovered genotypes with zero predicted peptide-HLA hits for each SARS-CoV-2 subunit. Columns indicate genotype frequency in each population. See `UncoveredGenotypes.xlsx`.

## Notes

https://github.com/gifford-lab/optivax

## References

Belgian Biosafety Server (2020). A randomized, double-blind, placebo-controlled Phase 1/2a study to evaluate the safety, reactogenicity, and immunogenicity of Ad26COVS1 in adults aged 18 to 65 years, inclusive and adults aged 65 years and older. https://www.biosafety.be/content/randomized-double-blind-placebo-controlled-phase-12a-study-evaluate-safety-reactogenicity. Accessed August 4, 2020.

BioNTech (2020). Pfizer and BioNTech Choose Lead mRNA Vaccine Candidate Against COVID-19 and Commence Pivotal Phase 2/3 Global Study. https://investors.biontech.de/news-releases/news-release-details/pfizer-and-biontech-choose-lead-mrna-vaccine-candidate-against. Accessed August 4, 2020.

Cohen, J. (2020). Merck, one of Big Pharma’s biggest players, reveals its COVID-19 vaccine and therapy plans. Science https://www.sciencemag.org/news/2020/05/merck-one-big-pharma-s-biggest-players-reveals-its-covid-19-vaccine-and-therapy-plans. Accessed August 4, 2020.

Dai, L., Zheng, T., Xu, K., Han, Y., Xu, L., Huang, E., An, Y., Cheng, Y., Li, S., Liu, M., et al. (2020). A universal design of betacoronavirus vaccines against COVID-19, MERS, and SARS. Cell 182, 722–733.

Dowdy, S.F. (2017). Overcoming cellular barriers for RNA therapeutics. Nature Biotechnology 35, 222–229.

Elbe, S., Buckland-Merrett, G. (2017). Data, disease and diplomacy: GISAID’s innovative contribution to global health. Global Challenges 1, 33–46.

Folegatti, P.M., Ewer, K.J., Aley, P.K., Angus, B., Becker, S., Belij-Rammerstorfer, S., Bellamy, D., Bibi, S., Bittaye, M., Clutterbuck, E.A., et al. (2020). Safety and immunogenicity of the ChAdOx1 nCoV-19 vaccine against SARS-CoV-2: a preliminary report of a phase 1/2, single-blind, randomised controlled trial. The Lancet 396, 467–478.

Gragert, L., Madbouly, A., Freeman, J., Maiers, M. (2013). Six-locus high resolution HLA haplotype frequencies derived from mixed-resolution DNA typing for the entire us donor registry. Human Immunology 74, 1313–1320.

Gupta, R., Jung, E., Brunak, S. (2004). Prediction of N-glycosylation sites in human proteins. In preparation URL: http://www.cbs.dtu.dk/services/NetNGlyc/.

Hadfield, J., Megill, C., Bell, S.M., Huddleston, J., Potter, B., Callender, C., Sagulenko, P., Bedford, T., Neher, R.A. (2018). Nextstrain: real-time tracking of pathogen evolution. Bioinformatics 34, 4121–4123.

Herst, C.V., Burkholz, S., Sidney, J., Sette, A., Harris, P.E., Massey, S., Brasel, T., Cunha-Neto, E., Rosa, D.S., Chao, W.C.H., et al. (2020). An effective CTL peptide vaccine for ebola zaire based on survivors’ CD8+ targeting of a particular nucleocapsid protein epitope with potential implications for COVID-19 vaccine design. Vaccine 38, 4464–4475.

Jackson, L.A., Anderson, E.J., Rouphael, N.G., Roberts, P.C., Makhene, M., Coler, R.N., McCullough, M.P., Chappell, J.D., Denison, M.R., Stevens, L.J., et al. (2020). An mRNA vaccine against SARS-CoV-2 —– preliminary report. New England Journal of Medicine https://doi.org/10.1056/NEJMoa2022483.

Jensen, K.K., Andreatta, M., Marcatili, P., Buus, S., Greenbaum, J.A., Yan, Z., Sette, A., Peters, B., Nielsen, M. (2018). Improved methods for predicting peptide binding affinity to MHC class II molecules. Immunology 154, 394–406.

Jurtz, V., Paul, S., Andreatta, M., Marcatili, P., Peters, B., Nielsen, M. (2017). NetMHCpan-4.0: improved peptide–MHC class I interaction predictions integrating eluted ligand and peptide binding affinity data. The Journal of Immunology 199, 3360–3368.

Klinger, M., Pepin, F., Wilkins, J., Asbury, T., Wittkop, T., Zheng, J., Moorhead, M., Faham, M. (2015). Multiplex identification of antigen-specific T cell receptors using a combination of immune assays and immune receptor sequencing. PLoS One 10, e0141561.

Kreiter, S., Selmi, A., Diken, M., Sebastian, M., Osterloh, P., Schild, H., Huber, C., Türeci, Ö., Sahin, U. (2008). Increased antigen presentation efficiency by coupling antigens to MHC class I trafficking signals. The Journal of Immunology 180, 309–318.

Liu, G., Carter, B., Bricken, T., Jain, S., Viard, M., Carrington, M., Gifford, D.K. (2020). Computationally optimized SARS-CoV-2 MHC class I and II vaccine formulations predicted to target human haplotype distributions. Cell Systems 11, 131–144.

Moyle, P.M., Toth, I. (2013). Modern subunit vaccines: development, components, and research opportunities. ChemMedChem 8, 360–376.

Mulligan, M.J., Lyke, K.E., Kitchin, N., Absalon, J., Gurtman, A., Lockhart, S., Neuzil, K., Raabe, V., Bailey, R., Swanson, K.A., et al. (2020). Phase 1/2 study of COVID-19 RNA vaccine BNT162b1 in adults. Nature, https://doi.org/10.1038/s41586-020-2639-4.

O’Donnell, T.J., Rubinsteyn, A., Bonsack, M., Riemer, A.B., Laserson, U., Hammerbacher, J. (2018). MHCflurry: open-source class I MHC binding affinity prediction. Cell Systems 7, 129–132.

O’Donnell, T.J., Rubinsteyn, A., Laserson, U. (2020). MHCflurry 2.0: Improved pan-allele prediction of MHC class I-presented peptides by incorporating antigen processing. Cell Systems 11, 42–48.

Pedregosa, F., Varoquaux, G., Gramfort, A., Michel, V., Thirion, B., Grisel, O., Blondel, M., Prettenhofer, P., Weiss, R., Dubourg, V., Vanderplas, J., Passos, A., Cournapeau, D., Brucher, M., Perrot, M., Duchesnay, E. (2011). Scikit-learn: Machine learning in Python. Journal of Machine Learning Research 12, 2825–2830.

Ramsuran, V., Naranbhai, V., Horowitz, A., Qi, Y., Martin, M.P., Yuki, Y., Gao, X., Walker-Sperling, V., Del Prete, G.Q., Schneider, D.K., et al. (2018). Elevated HLA-A expression impairs HIV control through inhibition of NKG2A-expressing cells. Science 359, 86–90.

Reynisson, B., Alvarez, B., Paul, S., Peters, B., Nielsen, M. (2020a). NetMHCpan-4.1 and NetMHCIIpan-4.0: improved predictions of MHC antigen presentation by concurrent motif deconvolution and integration of MS MHC eluted ligand data. Nucleic Acids Research 48, W449–W454.

Reynisson, B., Barra, C., Kaabinejadian, S., Hildebrand, W.H., Peters, B., Nielsen, M. (2020b). Improved prediction of MHC II antigen presentation through integration and motif deconvolution of mass spectrometry MHC eluted ligand data. Journal of Proteome Research 19, 2304–2315.

Sahin, U., Derhovanessian, E., Miller, M., Kloke, B.P., Simon, P., Löwer, M., Bukur, V., Tadmor, A.D., Luxemburger, U., Schrörs, B., et al. (2017). Personalized RNA mutanome vaccines mobilize poly-specific therapeutic immunity against cancer. Nature 547, 222–226.

Sahin, U., Muik, A., Derhovanessian, E., Vogler, I., Kranz, L.M., Vormehr, M., Baum, A., Pascal, K., Quandt, J., Maurus, D., et al. (2020). COVID-19 vaccine BNT162b1 elicits human antibody and TH1 T-cell responses. Nature, https://doi.org/10.1038/s41586-020-2814-7.

Sekine, T., Perez-Potti, A., Rivera-Ballesteros, O., Strålin, K., Gorin, J.B., Olsson, A., Llewellyn-Lacey, S., Kamal, H., Bogdanovic, G., Muschiol, S., et al. (2020). Robust T cell immunity in convalescent individuals with asymptomatic or mild COVID-19. Cell 183, 158–168.

Snyder, T.M., Gittelman, R.M., Klinger, M., May, D.H., Osborne, E.J., Taniguchi, R., Zahid, H.J., Kaplan, I.M., Dines, J.N., Noakes, M.N., et al. (2020). Magnitude and dynamics of the T-Cell response to SARS-CoV-2 infection at both individual and population levels. medRxiv, https://doi.org/10.1101/2020.07.31.20165647.

Srinivasan, S., Cui, H., Gao, Z., Liu, M., Lu, S., Mkandawire, W., Narykov, O., Sun, M., Korkin, D. (2020). Structural genomics of SARS-CoV-2 indicates evolutionary conserved functional regions of viral proteins. Viruses 12, 360.

Tang, M., Lautenberger, J.A., Gao, X., Sezgin, E., Hendrickson, S.L., Troyer, J.L., David, V.A., Guan, L., Mcintosh, C.E., Guo, X., et al. (2012). The principal genetic determinants for nasopharyngeal carcinoma in China involve the HLA class I antigen recognition groove. PLoS Genetics 8, e1003103.

Tian, J.H., Patel, N., Haupt, R., Zhou, H., Weston, S., Hammond, H., Lague, J., Portnoff, A.D., Norton, J., Guebre-Xabier, M., et al. (2020). SARS-CoV-2 spike glycoprotein vaccine candidate NVX-CoV2373 elicits immunogenicity in baboons and protection in mice. bioRxiv, https://doi.org/10.1101/2020.06.29.178509.

Virtanen, P., Gommers, R., Oliphant, T.E., Haberland, M., Reddy, T., Cournapeau, D., Burovski, E., Peterson, P., Weckesser, W., Bright, J., et al. (2020). SciPy 1.0: fundamental algorithms for scientific computing in python. Nature methods 17, 261–272.

Wang, N., Shang, J., Jiang, S., Du, L. (2020). Subunit vaccines against emerging pathogenic human coronaviruses. Frontiers in Microbiology 11, 298.

Wolff, J.A., Malone, R.W., Williams, P., Chong, W., Acsadi, G., Jani, A., Felgner, P.L. (1990). Direct gene transfer into mouse muscle in vivo. Science 247, 1465–1468.

World Health Organization (2020). Draft landscape of COVID-19 candidate vaccines. https://www.who.int/publications/m/item/draft-landscape-of-covid-19-candidate-vaccines. Accessed August 3, 2020.

Yu, J., Tostanoski, L.H., Peter, L., Mercado, N.B., McMahan, K., Mahrokhian, S.H., Nkolola, J.P., Liu, J., Li, Z., Chandrashekar, A., Martinez, D.R., Loos, C., Atyeo, C., Fischinger, S., Burke, J.S., Slein, M.D., Chen, Y., Zuiani, A., Lelis, F.J.N., Travers, M., Habibi, S., Pessaint, L., Van Ry, A., Blade, K., Brown, R., Cook, A., Finneyfrock, B., Dodson, A., Teow, E., Velasco, J., Zahn, R., Wegmann, F., Bondzie, E.A., Dagotto, G., Gebre, M.S., He, X., Jacob-Dolan, C., Kirilova, M., Kordana, N., Lin, Z., Maxfield, L.F., Nampanya, F., Nityanandam, R., Ventura, J.D., Wan, H., Cai, Y., Chen, B., Schmidt, A.G., Wesemann, D.R., Baric, R.S., Alter, G., Andersen, H., Lewis, M.G., Barouch, D.H. (2020). DNA vaccine protection against SARS-CoV-2 in rhesus macaques. Science 369, 806–811.

Zeng, H., Gifford, D.K. (2019). Quantification of uncertainty in peptide-MHC binding prediction improves high-affinity peptide selection for therapeutic design. Cell Systems 9, 159–166.

